# Competition-cooperation in the chemoautotrophic ecosystem of Movile Cave – first metagenomic approach on sediments

**DOI:** 10.1101/2022.05.19.492637

**Authors:** Iulia Chiciudean, Giancarlo Russo, Diana Felicia Bogdan, Erika Andrea Levei, Luchiana Faur, Alexandra Hillebrand-Voiculescu, Oana Teodora Moldovan, Horia Leonard Banciu

## Abstract

**Background:** Movile Cave (Dobrogea, SE Romania) hosts a subterranean chemoautotrophically-based ecosystem supported by a sulfidic thermal aquifer analogous to the deep-sea hydrothermal ecosystems. Our current understanding of Movile Cave microbiology has been confined to the thermal water proximity (no more than 2 m distant), with most studies focusing on the water-floating mat, which likely acts as the primary production powerhouse in this sulfidic ecosystem. To gain more insightful information on the functioning of the sulfidic Movile Cave ecosystem, we employed a metagenomics-resolved approach to reveal the microbiome diversity, metabolic potential, and interactions and infer its roles within the food webs in the sediments beyond the sulfidic thermal waters.

**Results:** A customized bioinformatics pipeline led to the recovery of 106 high-quality metagenome-assembled genomes from 7 cave sediment metagenomes. Assemblies’ taxonomy spanned 19 bacterial and three archaeal phyla with *Acidobacteriota, Chloroflexota, Proteobacteria, Planctomycetota, Ca*. Patescibacteria, *Thermoproteota, Methylomirabilota*, and *Ca*. Zixibacteria as prevalent phyla. Functional gene analyses allowed prediction of CO_2_ fixation, methanotrophy, sulfur and ammonia oxidation as possibly occurring in the explored sediments. Species Metabolic Coupling Analysis of metagenome-scale metabolic models revealed the highest competition-cooperation interactions in the sediments collected at the farthest distance from the sulfidic water. As a result of simulated metabolic interactions, autotrophs and methanotrophs were hypothesized as major donors of exchanged metabolites in the sediment communities. Cross-feeding dependencies were assumed only towards ‘currency’ molecules and inorganic compounds (O_2_, PO_4_^3-^, H^+^, Fe^2+^, Cu^2+^) in the sediment nearby sulfidic water, whereas hydrogen sulfide and methanol are predictably traded exclusively among communities dwelling in the distant gallery.

**Conclusions:** These findings suggest that the primary production potential of the Movile Cave expands way beyond its hydrothermal waters, enhancing our understanding of ecological interactions inside chemolithoautotrophically based subterranean ecosystems and their functioning.

## Background

Chemoautotrophically-based ecosystems known so far comprise deep sea hydrothermal vents and cold seeps [1–4], reduced sulfur-rich water and sediments of circumneutral saline and soda lakes [5–7] and sulfidic cave [8–10]. Only a handful of sulfidic cave ecosystems that rely totally or partially on chemosynthesis are explored to date, including Ayyalon Cave (Israel), Movile Cave (Romania), Frasassi Cave (Italy), Lower Kane Cave and Cesspool Cave (USA), and Villa Luz Cave (Mexico) [11–14, 9, 15, 16]. In these caves, organic matter is produced *in situ* by microorganisms that use inorganic compounds as nutrient and energy sources. Movile Cave (SE Romania) has been the first remarkable mentioning of a cave system defying the conventional view of subterranean ecosystems as being supported by aboveground photosynthetic productivity. Movile Cave is a small-surfaced, closed chemoautotrophic system [8] driven by in-house sulfur and methane oxidation and CO_2_ fixation as primary production processes. This cave, located in south-eastern Romania (Dobrogea region), developed in oolitic and fossil-rich limestone of Sarmatian age (Late Miocene) and sealed off during the Quaternary by a thick and impermeable layer of clays and loess [17]. Movile Cave has a complex geological evolution with an ongoing speleogenesis driven by two main processes: the sulfuric acid corrosion in the partially submerged, lower cave level; and the condensation-corrosion processes active in the upper level of the cave [18]. The upper gallery (approx. 200 m long) is dry whereas the lower gallery (approx. 40 m long) is partially flooded by sulfidic hydrothermal waters (T=∼20.7°C). The two levels converge in the so-called Lake Room (Fig. 1). Air pockets (Air Bells) are present in the submerged gallery (Fig. 1). Here, an active redox interface is created on both the water’s surface and the walls of the cave, which are colonized by floating microbial mats and biofilms. Most of our understanding of this ecosystem comes from studies on trophic chain components such as microbial mats associated with the hydrothermal water surface and its close proximity [19–22]. In addition, the rich endemic invertebrate fauna of Movile Cave gravitates in the proximity of the sulfidic water, being most abundant in the Air-Bells and Lake Room [15, 19] where the redox potential is higher, particularly in the Air-Bells water surface [23]. Based on these observations, it can be hypothesized that the chemoautotrophy potential of the ecosystem decreases with the distance from the hydrothermal waters, with the gradual emergence of two different niches: one based on chemoautotrophy and another, more like a regular, oligotrophic cave. Except from a snapshot investigation of the surface of rocks collected at about 2 m from the water surface [24], which revealed the presence of sulfur oxidizers and methylotrophic bacteria, Movile Cave has not yet been explored for the microbial life associated with sediments distant from the sulfidic water. The aim of this study is to address the diversity and metabolic potential of microbial communities inhabiting the sediments beyond the hydrothermal waters of Movile Cave. We performed metagenomic analyses and genome-resolved inferences of ecological interactions to examine the taxonomy, abundance, metabolic features, and interactions of sediment microbial communities to understand their role within the complex food web of Movile Cave with 52 endemic invertebrate species [6].

**Figure 1.**
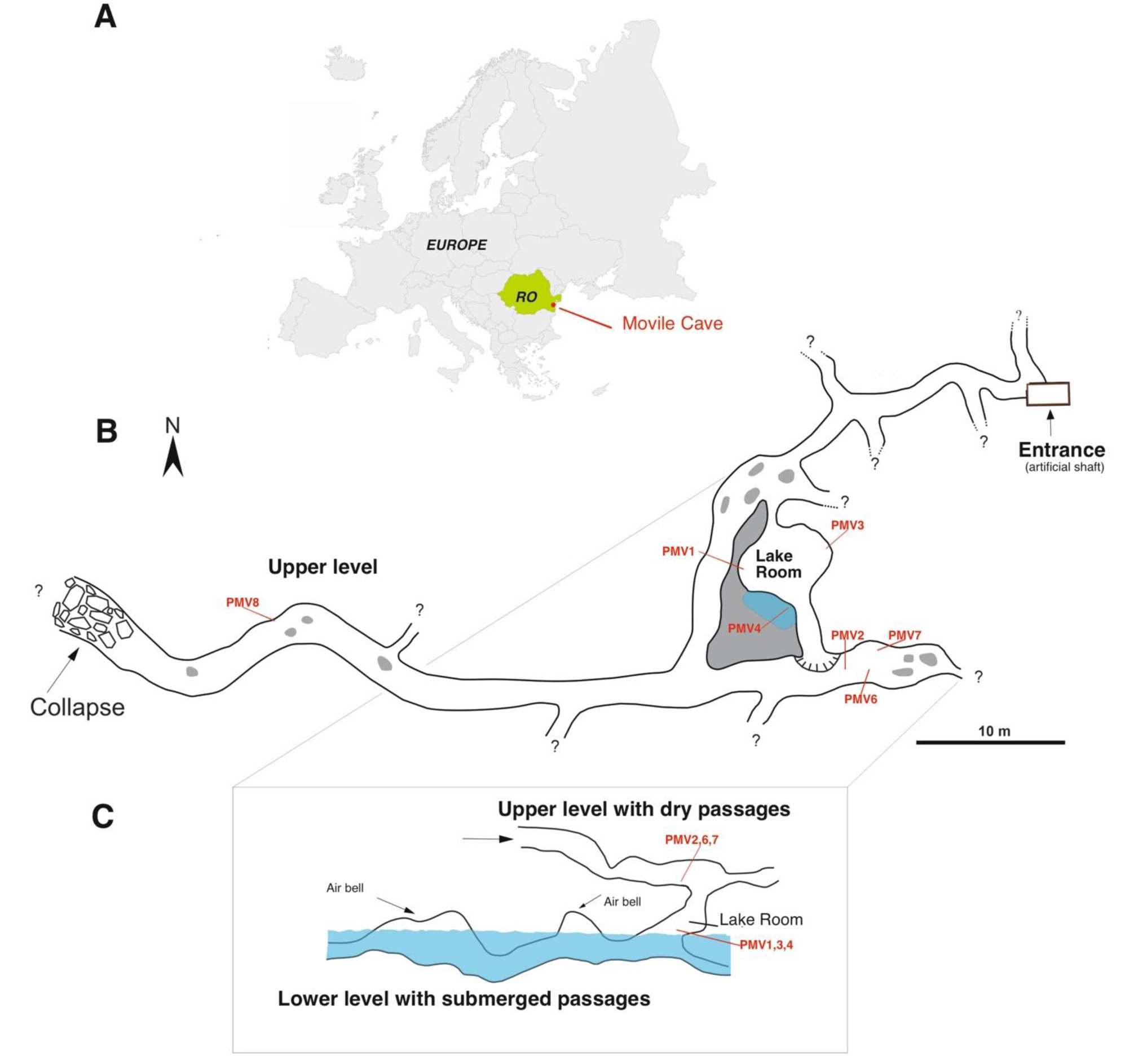
Schematic representations of Movile Cave map (B) and profile (C) with its localization in Romania and Europe (A). The sampling sites from this study are PMV1-PMV8 (modified from Sarbu et al., 2019 [25])

## Materials and methods

### Study site and sample collection

Sediment samples were collected from seven sites located in the upper and lower levels (galleries) of the cave (Fig. 1) in December 2019. Sampling sites were selected based on their physical characteristics (explained in detail in Table 1 and Additional file 1: Figure S1) and position related to the Lake Room. In brief, three sampling sites were near the water, in the Lake Room (PMV 1, PMV3, PMV4), and the other four (PMV2, PMV6, PMV7, PMV8) in the upper, dry gallery. As an observation, PMV5 sample consisted of sulfidic water and was not the subject of this study. Stable environmental conditions typical to a subterranean habitat combined with hydrothermal activity drive a constant temperature of the water and air at ≈20.7°C within Movile Cave. Aseptically collected top sediments (up to 50 g) were kept at 4°C after sampling and while transporting, then at -20°C until processing.

**Table 1.** Description of the sites sampled in Movile Cave during December 2019. Stations in the lower level are marked in blue, while stations in the upper level are in brown.

| Sample codes | Sampling sites characteristics | Sediment description | Microclimate |
| --- | --- | --- | --- |
| PMV1 | Lower gallery, Lake Room, inside a wall diverticulum | Fine grey silt | Air: Humidity=99%; Temp=$20.8^{\circ}\text{C}$; $\text{CO}_2=17,420$ ppm |
| PMV2 | Upper gallery; floor, 50-70 cm away from the wall | Yellowish silt/clay | Air: Humidity=98%; T=$20.6^{\circ}\text{C}$; $\text{CO}_2=17,010$ ppm |
| PMV3 | Lower gallery, Lake Room depressions in the wall | Fine grey silt | Air: Humidity=99%; Temp=$20.8^{\circ}\text{C}$; $\text{CO}_2=17,420$ ppm |
| PMV4 | Lower gallery, Lake Room; water-sediment interface (water edge) | Fine black silt | Water: pH=7.26; $\text{O}_2=0.12$ mg/L; EC=1510 $\mu\text{S}$; TDS=754 ppm; Temp=$21.5^{\circ}\text{C}$<br>Air: Humidity=99%; Temp=$20.8^{\circ}\text{C}$; $\text{CO}_2=17,420$ ppm |
| PMV6 | Upper gallery; floor, 50-70 cm away from the left wall | Coarse yellowish silt/clay | Air: Humidity=98%; T=$20.6^{\circ}\text{C}$; $\text{CO}_2=17,010$ ppm |
| PMV7 | Upper gallery; floor, 50-70 cm away from the right wall | Yellowish silt/clay | Air: Humidity=98%; T=$20.6^{\circ}\text{C}$; $\text{CO}_2=17,010$ ppm |
| PMV8 | Upper gallery; wall diverticulum. | Coarse grey silt | Air: Humidity=97%; T=$20.8^{\circ}\text{C}$; $\text{CO}_2=14.800$ ppm |

### Mineralogy and geochemistry measurements

Powdered X-ray diffraction analyses were performed on sediments in order to establish their mineralogy. Samples were analyzed with a Rigaku Ultima IV diffractometer in parallel beam geometry equipped with CuKα radiation (wavelength 1.5406 Å). The XRD patterns were collected in 2Θ range between 5 to 80 with a speed of 2°/min and a step size of 0.02°. PDXL software from Rigaku, connected to the ICDD database was used for phase identification. The quantitative determination was made using the RIR (Reference Intensity Ratio) methodology.

The pH and electric conductivity were measured in 1:5 solid to water extract using a Seven Excellence (Mettler Toledo) multimeter. The N, C and H were measured on freeze-dried samples using a Flash 2000 (Thermo Scientific, Thermo Fisher Scientific, Waltham, MA, United States) analyzer. For measurement of metals, samples were digested using a 1:3 mixture of 65 % HNO_3_ and 37 % HCl at reflux conditions. Major elements (Na, K, Ca, Mg, Al, Fe, S, P) in the digested samples were determined by inductively coupled plasma optical emission spectrometry (ICP-OES) using an 5300 Optima DV spectrometer (Perkin Elmer, Shelton, CT, United States), while trace metals and metalloids by inductively coupled plasma mass-spectrometry (ICP-MS) using an ELAN DRC II spectrometer (Perkin Elmer, Waltham, MA, United States). Additional measurements were done for the aqueous fraction of the PMV4 sample. Total nitrogen (TN) was measured in unfiltered water while dissolved nitrogen (DN) was measured in water samples filtered through 0.45 µm pore size PTFE syringe filters by catalytic combustion and chemiluminescence detection, using a Multi N/C 2100 (Analytik Jena, Jena, Germany) analyzer. Total carbon (TC) and total inorganic carbon (TIC) were measured from unfiltered samples while dissolved carbon (DC) and dissolved inorganic carbon (DIC) from water samples filtered through 0.45 µm pore size PTFE syringe filters by catalytic combustion and non-dispersive infrared detection, using a Multi N/C 2100 (Analytik Jena, Jena, Germany) analyzer. Total dissolved solids (TDS) were determined by gravimetric method. Anions (NO_2_^-^, NO_3_^-^, F^-^, Cl^-^, PO_4_^3-^, SO_4_^2-^) were measured by ion-chromatography, using a 761 IC compact ion chromatograph (Metrohm, Herisau, Switzerland), equipped with a Metrosep 5–100/4 column and a Metrosep A Supp 4/5 mm guard column after filtration of samples through 0.45 μm cellulose acetate membrane filters. Na, Mg, K and Ca were measured after in the filtered samples after acidulation with 65 % HNO_3_ to a pH<2, by ICP-OES.

### DNA extraction and sequencing

Community DNA from sediments was extracted using the DNeasyPowerMax Soil Kit (Qiagen, Gaithersburg, MD, United States); precipitated with 1:10 (v:v) of 3 M Na-Acetate (pH 5.2) and 1:1 (v:v) of 100 % ethanol; air-dried and finally resuspended in 100 µL of sterile TE buffer. DNA extracts were quantified with Qubit™dsDNA BR Assay Kit and a Qubit 4.0 fluorometer (Invitrogen, Thermo Fisher Scientific, Waltham, MA, United States). Novogene Europe Co., Ltd (Cambridge, UK) performed the library preparation and shotgun metagenomics sequencing. For library preparation Illumina protocol NEBNext® Ultra™DNA Library Prep Kit (NEB, USA) was used starting from 200 ng DNA. Barcodes were added to each sample. Sonication was used to fragment the DNA to a size of around 350 bp. The PCR amplification was conducted after the DNA fragments were end-polished, A-tailed, and ligated with the adapter. AMPure XP system was used for PCR product purification. Agilent 2100 Bioanalyzer was used to analyze the size distribution of the DNA library, and real-time PCR was performed to quantify DNA libraries. The samples were sequenced and paired-end reads were generated (PE150) on the Illumina NovaSeq 6000 platform.

### Sequencing, assembly, and read annotation

Metagenomic raw reads (fastq) were quality-filtered and adapter-trimmed using Trimmomatic v0.39 [26] and FastQC v0.11.9 [27]. The resulting sequences were assembled into contigs using MEGAHIT v1.2.9 [28], with the following parameters: no-mercy, k-min = 31, k-max = 101, k-step = 101. Assembled contigs (> 1 kb) were binned using two binners: MaxBin v2.0 [29] and MetaBAT2 v2.12.1 [30]. DAS Tool (v1.1.2) was applied to integrate the bins generated from the two methods [31]. Only bins with over 70% completion and less than 10% of contamination according to CheckM v1.1.3 [32] were analyzed. Protein-coding sequences (CDS) of selected metagenome-assembled genomes (MAGs) were predicted and annotated with Prodigal v2.6.1 [33] and DIAMOND v2.0.5 [34] against Uniprot databases. Furthermore, the functional annotation of CDS and the prediction of metabolic pathways related to sulfur, nitrogen, carbon, and methane metabolism was carried out using the KEGG’s Ghost Koala tool [35]. For confirmation, CDS of interest were identified by blastp against NCBI nr database.

### Taxonomic assignment and abundance of MAGs

Taxonomic classification of MAGs was assessed using Genome Taxonomy Database Toolkit (GTDB-Tk v1.5.0, reference data for GTDB R06-RS202) [36] based on 120 bacterial and 122 archaeal marker genes. For phylogenetic analysis, closely related MAGs, reference genomes, or type strains genomes were selected from GTDB (July 2021). The marker genes alignment was concatenated and then imported into R using the *readAA Multiple Alignment* function from the package *Biostrings*. Phylogenetic distances were calculated using the function *dist*.*alignment* from the package *seqinr*, and a phylogenetic tree was built using the neighbor-joining tree estimation as implemented in the function *nj* in the R package *ape*. The tree was visualized using the online iTOL tool [37].

Finally, for MAGs coverage, the filtered shotgun reads were mapped back to contigs belonging to each MAG with Minimap2 v2.21 [38], and the coverage was summarized with the *coverage* command from SAMtools v1.11 [39]. The relative abundance of each MAG in each metagenome was given by the number of reads mapped per Kb of MAG divided by Gb of corresponding metagenome (RPKG)[40].

### Genome-scale metabolic reconstruction and interaction models

Genome-scale metabolic models derived from MAGs (metaGEMs) were reconstructed using CarveMev. 1.5.1 [41], with the default CPLEX solver (v. 12.8.0.0) (IBM,2019). Two sets of metaGEMs were reconstructed: one set without gap-filling for any particular medium, based only on genetic evidence, and another set gap-filled for a custom minimal medium that would guarantee the models’ growth in a nutrient-poor environment as found in Movile Cave. MEMOTE tests suite [42] was applied on the gap-fill models initialized for the minimal medium to ensure that the metaGEMs could generate biomass and reproduce growth in a scarce environment. The custom minimal medium composition is detailed in Additional file 2: Table S1. Community simulations were performed with SMETANA 1.1.0 [43] using the reconstructed metaGEMs. For the simulations, the samples were divided into two groups based on their distance from the sulfidic water, namely samples from the lower gallery, in the Lake Room (PMV1, PMV3, PMV4) and samples from the upper gallery (PMV2, PMV6, PMV7, PMV8). Metabolic interactions were assessed for each sample, and metabolic dependencies were calculated for each condition (lower vs. upper gallery). Community global interactions SMETANA algorithms MRO (*<u>m</u>*etabolic *<u>r</u>*esource *<u>o</u>*verlap) and MIR (*<u>m</u>*etabolic *i*nteraction *<u>p</u>*otential) were applied for both sets of reconstructed metaGEMs assuming complete or minimal media. MRO/MIR scores were used as a measure of how much the species compete for the same metabolites (MRO) and as a measure of how many metabolites the species can share to decrease their dependency on external resources (MIP). Only the gap-filled models were used for metabolic dependency, calculated assuming a complete medium. The complete medium composition relies on the BIGG Database [44] (http://bigg.ucsd.edu/). By doing so, we imposed the growth of community members on the minimal medium as a constraint but relaxed the community simulation to all possible metabolic exchanges of a complete environment (to identify the metabolic exchanges essential for the community’s survival). SMETANA scores estimate the strength of metabolic coupling (metabolites exchange) in a community. Only a SMETANA score of 1 was considered for analysis. The interactions characterized by a SMETANA score of 1 are considered essential metabolic exchange (essentialities) for the community, and this metabolic coupling is considered a “dependency”. SMETANA score was used to highlight the cross-feeding interactions and the keystone MAGs (phyla). Significantly different (p < 0.001) compound exchanges across conditions were identified by the Wilcoxon rank-sum test using *compare_means* function of *ggpubr*R package. Models ‘ reconstruction and community simulations parameters, variants and the simulation codes are summarized and detailed in Additional file 2: Table S2, S3.

The figures were generated using the R packages *ggplot2* [45], *pheatmap* [46], and *circlize* [47].

### Statistical analysis

We used the multidimensional scaling (MDS) to represent the sediment samples in a bi-dimensional space. For the MDS we used a proximity matrix of similarity between the objects (the chemical characteristics of the sediment samples) to the coordinates of these same objects in a 2-dimensional space so that the objects may be visualized easily. The analysis was done in XLSTAT 2021.4.1 (Addinsoft, New York).

## Results

### Physicochemical and mineralogical characterization

The sediments had a slightly alkaline pH, in the range of 7.9 - 9.2, and low (PMV1, PMV7 and PMV8) or high (PMV 2, PMV3, PMV4 and PMV6) electrical conductivity ranging from 60.6 to 1,738.8 µS cm^−1^ (Table 2). The physico-chemical characteristics of the sediment samples are shown in Table 2. Total carbon (TOC) and total nitrogen (TN) content, measured for the aqueous fraction of the sample collected from the water edge (PMV4) were low (TOC = 14.9 mg L^−1^, TN = 3.7 mg L^−1^) with a TOC/TN of ∼4 mg L^−1^ (Additional file 3: Table S4).

**Table 2.** Physico-chemical and mineralogical composition of the sediment samples in Movile Cave.

| **Parameter (unit)** | **PMV1** | **PMV2** | **PMV3** | **PMV4** | **PMV6** | **PMV7** | **PMV8** |
| --- | --- | --- | --- | --- | --- | --- | --- |
| **pH** | 8.5 | 8.3 | 7.9 | 8.3 | 8.1 | 8.9 | 9.2 |
| **Electrical conductivity** ($\mu\text{S cm}^{-1}$) | 270 | 751 | 955 | 1 237 | 1 739 | 119 | 60.6 |
| **N** (%SU) | 0.21 | <0.01 | <0.01 | 1.37 | <0.01 | <0.01 | <0.01 |
| **C** (%SU) | 9.1 | 4.8 | 3.9 | 8.8 | 8.0 | 6.6 | 4.1 |
| **H** (%SU) | 0.4 | 0.4 | 0.8 | 0.3 | 0.4 | 0.4 | 0.6 |
| **Ca** ($\text{mg kg}^{-1}$) | 172 900 | 125 000 | 119 100 | 255 100 | 159 500 | 136 900 | 92 300 |
| **Al** ($\text{mg kg}^{-1}$) | 21 450 | 25 950 | 19 520 | 12 635 | 11 630 | 29 540 | 38 490 |
| **Mg** ($\text{mg kg}^{-1}$) | 42 960 | 29 860 | 6 834 | 27 590 | 51 980 | 36 320 | 28 120 |
| **Fe** ($\text{mg kg}^{-1}$) | 10 160 | 10 540 | 21 470 | 10 030 | 5 545 | 10 140 | 12 780 |
| **K** ($\text{mg kg}^{-1}$) | 5 103 | 4 279 | 4 983 | 2 145 | 2 140 | 5 816 | 6 185 |
| **Na** ($\text{mg kg}^{-1}$) | 415 | 289 | 126 | 336 | 564 | 374 | 240 |
| **S** ($\text{mg kg}^{-1}$) | 902 | 3 003 | 13 270 | 5 730 | 30 480 | 1 610 | 136 |
| **P** ($\text{mg kg}^{-1}$) | 162 | 239 | 357 | 110 | 163 | 186 | 124 |
| **Mn** ($\text{mg kg}^{-1}$) | 278.0 | 216 | 138 | 38.5 | 99.1 | 184 | 343 |
| **Ba** ($\text{mg kg}^{-1}$) | 86.5 | 135 | 60.3 | 429 | 87.3 | 183 | 110 |
| **Ti** ($\text{mg kg}^{-1}$) | 202 | 246 | 127 | 230 | 149 | 196 | 87.8 |
| **Sr** ($\text{mg kg}^{-1}$) | 75.4 | 106 | 52.2 | 1060 | 270 | 91.6 | 48.2 |
| **V** ($\text{mg kg}^{-1}$) | 29.6 | 26.9 | 261 | 15.5 | 16.4 | 24.2 | 33.6 |
| **As** ($\text{mg kg}^{-1}$) | 39.1 | 24.7 | 322.5 | 20.6 | 24.8 | 26.4 | 19.5 |
| **Zn** ($\text{mg kg}^{-1}$) | 9.9 | 14.9 | 19.4 | 138 | 5.7 | 10.8 | 17.7 |
| **Cr** ($\text{mg kg}^{-1}$) | 14.9 | 20.4 | 101 | 8.0 | 4.7 | 11.9 | 11.1 |
| **Mineralogy**<br>(>5%) | dolomite,<br>quartz,<br>calcite | quartz,<br>calcite,<br>muscovite | calcite | calcite,<br>quartz,<br>dolomite,<br>muscovite | dolomite,<br>gypsum,<br>quartz | dolomite,<br>calcite,<br>quartz,<br>muscovite | dolomite,<br>muscovite,<br>aragonite |

In the upper cave gallery, the identified minerals are calcite, dolomite, aragonite, quartz, muscovite, gypsum, and in small amounts, goethite. Near the lake, in the lower gallery, the association is dominated by dolomite and calcite, followed by quartz, gypsum and muscovite (Table 2). Carbonates are represented mainly by calcite derived from the limestone bedrock, dolomite and aragonite. Dolomite is present in substantial amounts in both levels of the cave. The bedrock mineralogy reveals the presence of magnesium-rich carbonate, which could be the explanation for the occurrence of aragonite, a signature of PMV8 mineralogy. Samples mineralogy is shown in Table 2 and Additional file 3: Table S5.

A multidimensional scaling (MDS; Fig.2) analysis shown that PMV6 and PMV8 are more dissimilar from the other samples, PMV6 having the higher electrical conductivity and higher S, Na, Mg, Ca combined with low K, Al and Fe content. The presence of gypsum (sulfate mineral) in that sample (Table 2) explains the high content of S and Ca. In contrast, PMV8 had the lowest electrical conductivity and content in S, Ca (no gypsum in this sample), with a higher concentration of Al, Li, and Na. PMV3 and PMV4 were also separated, while PMV1, 2 and 7 were more similar along the MDS 1^st^ Dimension. PMV3 had low Na and Mg, and high Fe (the only sample where goethite - iron(III) oxide-hydroxide - is well represented), V, As, Cr, etc. PMV4 had the highest content of Ca, Ni, and Ba that probably gives the black color to the sample. The other samples (LMV1, 2, and 7) were less dissimilar. All three had muscovite (hydrated phyllosilicate mineral of aluminum and potassium) and quartz (SiO_2_) in their mineralogical composition.

**Fig. 2.**
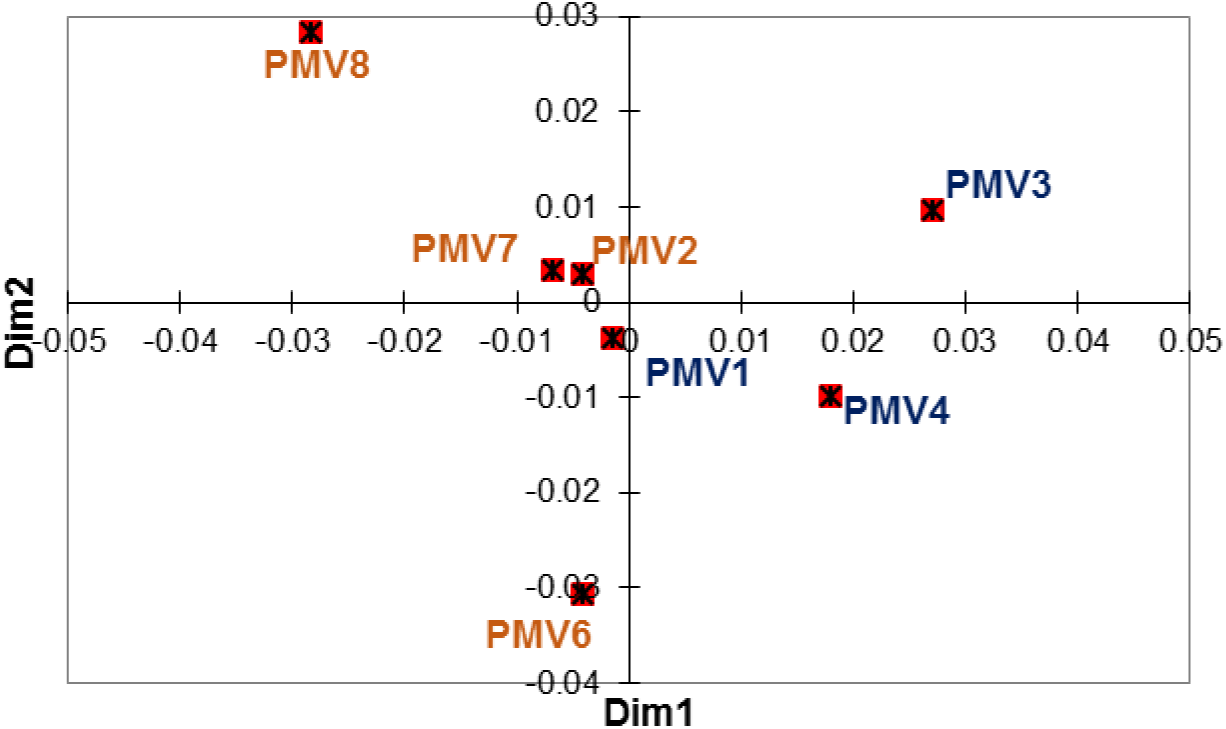
The plot of MDS analysis based on the chemical composition of sediments sampled from Movile Cave. The analysis shows the strong separation of PMV8 and PMV6 samples and a less strong separation of PMV3 and PMV4 from the other sediment samples. The Kruskal’s stress (1) = 0.207.

### MAGs recovery and phylogeny

A total of 106 medium- to high-quality MAGs (> 70% complete and < 10% contamination) [48] were recovered from the sediment metagenomes of Movile Cave. The taxonomic assignment performed by using GTDB-tk indicated that the recovered MAGs span 19 bac terial and three archaeal phyla (Fig. 3C.). Out of those, 4 were candidate phyla *Ca*.Patescibacteria, KSB1, Krumholzibacteriota and Zixibacteria. The percentages of reads mapped to MAGs varied among samples from ca. 80 % to 13 % for MAG recovery efficiency. Based on the relative abundance of MAGs in the sample (expressed as RPKG), the most abundant phyla were *Acidobacteriota, Chloroflexota, Proteobacteria* and *Planctomycetota* (>20 RPKGs in any of the samples) (Fig. 3A.). For the lower gallery, *Chloroflexota* was abundant in PMV1 and PMV3, while *Proteobacteria* (class *Gammaproteobacteria*) dominated PMV4. In the upper, dry gallery, *Acidobacteriota* was the most abundant phylum for PMV8 and PMV7. For the phyla with a lower relative abundance (< 20 RPKGs), *Ca*.Patescibacteria was the signature phylum for PMV2, *Thermoproteota* for PMV3, *Methylomirabilota* for PMV7 and *Ca*.Zixibacteria for PMV8 (Fig. 3B.).

**Figure 3.**
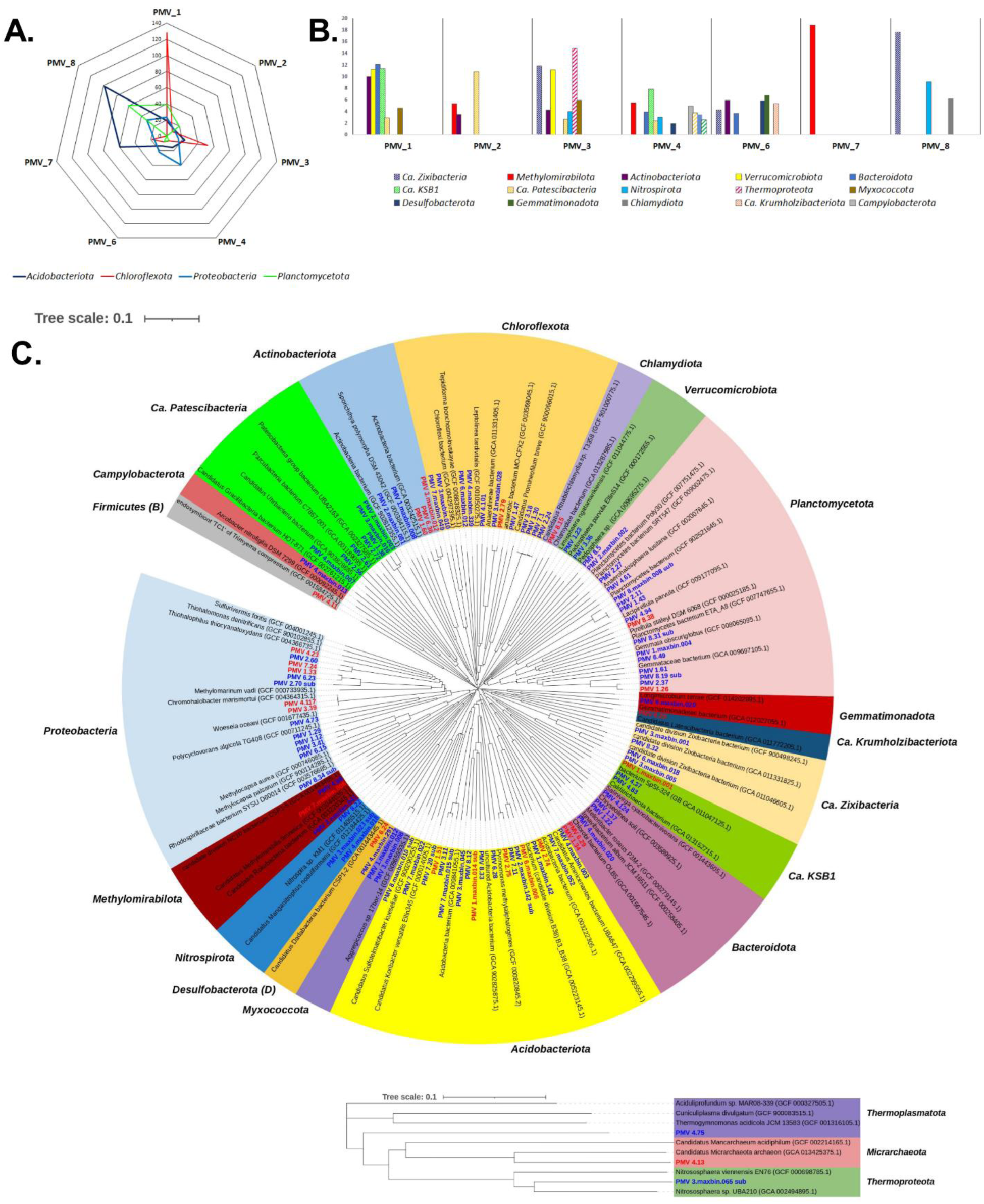
The abundance and phylogeny of MAGs recovered from sediments of Movile Cave. **A**. Most abundant phyla (relative abundance >20 RPKGs). **B**. Less abundant phyla (relative abundance < 20 RPKGs). **C**. Phylogenetic tree of MAGs from Movile Cave sediments including their closely related MAGs from GTDB (GCA) and NCBI type material genomes (GCF) (type strain and/or reference genomes). The neighbor-joining phylogenetic tree was constructed based on the GTDB marker genes. The MAGs detected in this study are shown in *<u>blue</u>* or *<u>red</u>* for medium- or high quality MAGs, respectively.

Out of the 106 MAGs, 25 were classified as high-quality (> 90% complete and < 5% contamination) (Fig. 3C.), but only two were assigned to species level (PMV3.39 and PMV4.117 affiliated to *Chromohalobacter marismortui*) with available genomes in the NCBI database (gANI> 95%). One-third of the MAGs (32 of 106) was classified by RED (Relative Evolutionary Divergence) as new species of established GTDB taxa [49]. Given the taxonomic novelty, RefSeq type material genomes (GCF) and representatives MAGs from GeneBank (GCA) were incorporated in the MAGs phylogenetic tree construction for a better representation (Fig. 3C.)

MAGs attributes, extended taxonomy and abundance are summarized in Additional file 4: Table S6 to S8.

### Potential for biogeochemical cycling of S and N, CH_4_ oxidation and CO_2_ fixation

#### Sulfur metabolism

Microbial sulfur cycling has been proposed as a driving force for bacterial proliferation in microbial mats of Movile Cave [24], but not investigated in the cave’s sediments. Here, we examined the presence of the marker genes for sulfur cycle in the metagenomes of Movile Cave’s sediments. For **sulfur oxidation** (Fig. 4), the complete canonical pathway for thiosulfate (S_2_O_3_^2–^) to sulfate (SO_4_^2-^) conversion (*soxAX, B,YZ*, (*CD*)) was annotated in the water edge dataset PMV4; and partially annotated in the upper gallery datasets PMV2 (*soxZ)*, PMV6 (*sox*(*CD*), *Y)* and PMV7 (*soxYZ*). In the PMV4 dataset, Sox-complex was encoded in MAGs affiliated to order *Thiohalomonadales* (class *Gammaproteobacteria*) (PMV4.23) and family *Arcobacteraceae* (class *Campylobacteria*) (PMV4_maxbin.013). The *Thiohalomonadales*-affiliated MAG encoded an incomplete thiosulfate-oxidation pathway lacking Sox(CD) and carrying multiple copies of *soxB*. For the sulfide (HS^-^) oxidation, the MAG encoded both the pathway to elemental sulfur (S^0^) via flavocytochrome c sulfide dehydrogenase (*fccAB*), and to sulfate (SO_4_^2-^) via the reverse-operating dissimilatory sulfite reductase pathway (*dsrAB*+*aprAB*+*sat*). The complete path for sulfite (SO_3_^2-^) oxidation to sulfate (SO_4_^2-^) (*soeABC*) was also annotated here. This *Thiohalomonadales* MAG also encoded (*hydBD*) the production of HS^-^ from polysulfides by sulfhydrogenase complex (*hyd*(*G,B*),(*A,D*)).

**Figure 4.**
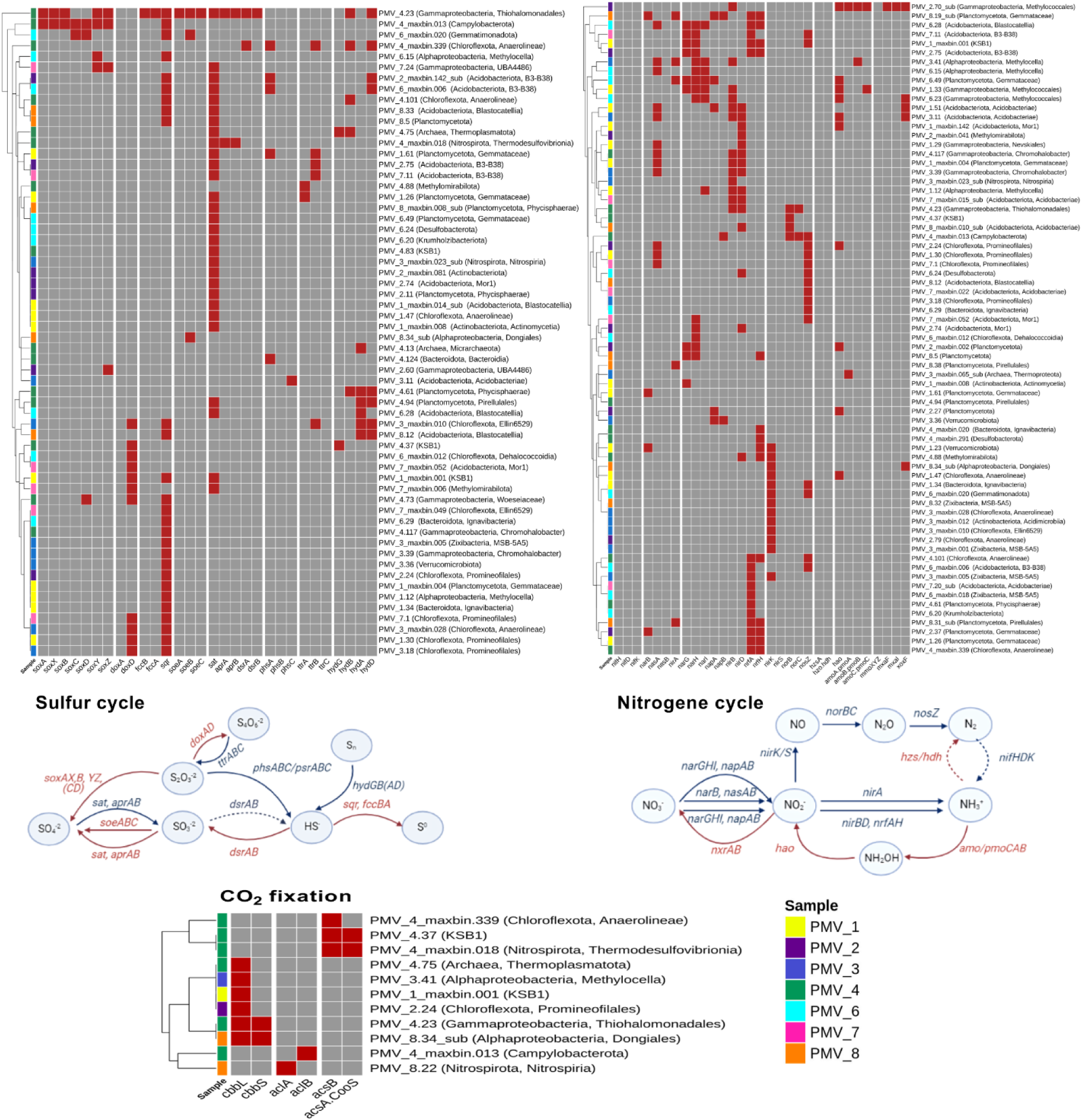
Overview of pathways and genes involved in sulfur, nitrogen cycling, methane oxidation and CO2 fixation encoded by MAGs recovered from Movile Cave sediments. The color scheme gives the presence/absence of functional genes: presence is indicated in red, absence in grey. The involvement of each gene in specific pathways is indicated in the diagrams. Red arrows indicate oxidation, and the blue arrows show the reduction of compounds. Full arrows indicate the enzymatic reactions for which the coding genes were found in the datasets based on the analyzed MAGs. The dotted arrows show enzymatic reactions absent in the datasets.

In the rest of the datasets, Sox(YZ) subunits were encoded in *Proteobacteria* MAGs and Sox(CD) in a *Gemmatimonadota* MAG (PMV6_maxbin.020). Thiosulfate oxidation with tetrathionate (S_4_O_6_ ^2-^) formation was partially encoded (*doxD*) in all datasets except for PMV8, and mostly in bins affiliated to *Chloroflexota*. Sulfide:quinone reductase (*sqr*) for the oxidation of HS^-^ with the zero-valent sulfur S^0^ formation was encoded in all datasets. The oxidation of sulfide via SQR was predicted as a widespread trait encoded in 27 MAGs. The pathway for sulfite (SO_3_ ^2-^) oxidation to sulfate (SO_4_ ^2-^) was also partially encoded (*soeB*) in PMV6 and PMV8 datasets. Sulfur respiration, dissimilatory **reduction** of oxidized sulfur compounds (SO_4_^2-^, SO_3_^2-^, S_2_O_3_^2–^, S^0^) coupled with sulfide (HS^-^) production, were also detected. Sulfate (SO_4_^2-^) to sulfite (SO_3_^2-^) reduction (*sat*+*aprAB*) was present in the PMV4 dataset, in a MAG affiliated to order *Thermodesulfovibrionales* (phylum *Nitrospirota*) (PMV4_maxbin.018). An NCBI blast (blastx against RefSeq Select database) showed the AprAB sequences similarities to *Deltaproteobacteria (*≈73% to 80%), placing them in the direct-operating dissimilatory pathway.

Thiosulfate/polysulfate (S_2_O_3_^2–^/S^0^) disproportionation and tetrathionate respiration were partially encoded (*phsA, phsC; ttrA/B*) in different phylogenetic groups, but a *Planctomycetota*-affiliated MAG (PMV1.61) that might encode both as *ttrB* and *phsA* were present in the bin. The production of hydrogen sulfide from polysulfides by sulfhydrogenase complex (*hyd*(*G,B*),(*A,D*)) was partially encoded especially by MAGs affiliated to *Acidobacteriota, Chloroflexota* and *Planctomycetota*.

#### Nitrogen metabolism

Analogous to sulfur, we addressed the potential for biogeochemical N cycling across sediments of Movile Cave. Primary producers capable to obtain energy for autotrophy by **nitrogen oxidation** (nitrification) were questioned. The first step of *nitrification*, the ammonia NH_3_^+^ to nitrite NO_2_^-^ oxidation was suggested in Movile Cave sediments as ammonia monooxygenase AMO (*amoABC*) was encoded in PMV1 to PMV3 datasets. All three subunits of AMO’s were encoded in an order *Methylococcales*-affiliated MAG (PMV2.70_sub). However, the *amoABC* sequences were homologous (>65% similarity) to the sequence encoding particulate methane monooxygenase pMMO in the genus *Methyloterricola*. The methanotrophs can oxidize both substrates (NH_3_ and CH_4_) but grow only on their characteristic substrate [50]. Interestingly, AmoA-like subunit was annotated in an archaeon MAG affiliated to *Nitrososphaera* genus (phylum *Thermoproteota*) (PMV3.maxbin.65_sub). For the second step of nitrification, the oxidation of NO_2_^-^ to NO_3_^-^ none of the bins encoded the nitrite oxidoreductase (*nxrAB*) found in known nitrite-oxidizing genera. Also, no marker genes (*hzsA* and *hzo*) for the anaerobic ammonia oxidation (*anammox*) (NH_3_ to N_2_H_4_ and then to N_2_) were annotated in the Movile datasets.

In the chemoautotrophic cave environment, the nitrogen demand can be met by converting inorganic nitrogen to a biologically useful form by **nitrogen reduction**. The conversion involves microbial *dinitrogen fixation* or *nitrate assimilation*. Surprisingly, the N_2_ fixation pathway was not detected in any sediment datasets, as no nitrogenase genes (*nif*) for the reduction of atmospheric molecular nitrogen (N_2_) to ammonia (NH_3_^+^) was annotated. The assimilatory process (NO_3_^-^, NO_2_^-^ conversion to NH_3_^+^), catalyzed by nitrate and nitrite reductases (*nasA/B, narB; nirA*), was fully encoded in PMV3, PMV6, and PMV8 datasets. The bins that encoded the nitrate-assimilatory enzymes were taxonomically diverse as the ability is however widely distributed among bacteria.

The first step in the *dissimilatory* nitrate reduction to ammonia (DNRA) and *denitrification*, NO_3_^-^ to NO_2_^-^ reduction, was fully encoded as the cytosolic nitrate reductases (*narGHI*) (PMV1, PMV6) or as the periplasmic nitrate reductases (*napAB*) (PMV3, PMV8). The NO_2_^-^ to NH_3_^+^step of DNRA was encoded in all datasets as the cytoplasmic nitrite reductase (*nirBD*) or the periplasmic nitrite reductase complex (*nrfAH*). MAGs that encoded at least partially both steps of the DNRA pathway were affiliated to *Acidobacteriota* (PMV1.51 (*Nap-Nir*); PMV2.75 (*Nar-Nrf*); PMV7.11 (*Nar-Nrf*)), *Planctomycetota* (PMV8.19_sub (*Nap-Nrf*); PMV8.5 (*Nar-Nrf*)) and *Gammaproteobacteria* (PMV6.23 (*Nar-Nir*)). Nitrite reductase (*nirK*), the hallmark enzyme of denitrification (NO_2_^-^ to (NO) conversion) was encoded in all datasets except PMV7, in bins taxonomically unrelated with common denitrifyers, including *Ca*.Zixibacteria (PMV8.32) and *Ca*.Methylomirabilis (PMV4.88). The conversion of NO to nitrous oxide (N_2_O) was fully encoded as nitric oxide reductase (*norBC*) in PMV4 in the sulfur-oxidizing *Thiohalomonadales* MAG (PMV4.23). The last step of denitrification, conversion of N_2_O to N_2_, carried out by an oxygen-sensitive enzyme, the nitrous-oxide reductase (*nosZ*), was annotated in all data sets and across taxonomically diverse bins.

#### Methane metabolism

Via **methane oxidation**, methanotrophs can metabolize methane as their source of carbon and energy. In the oxidation pathway, methane (CH_4_) is oxidized to methanol (CH_3_OH) and then to formaldehyde (CH_2_O) which is incorporated into organic compounds via the serine or the ribulose monophosphate (RuMP) pathway. As previously mentioned, all subunits of the particulate methane monooxygenase pMMO (*pmoABC*) were predicted in the order *Methylococcales* MAG (PMV2.70_sub). Separated subunits of pMMO were also annotated in MAGs affiliated to *Methylococcales* order (PMV1.33) and *Methylocella* genus (PMV 3.41). No soluble methane monooxygenase (sMMO) was annotated across investigated metagenomes. The methanol oxidation to formaldehyde appeared encoded as Ln^3+^-dependent methanol dehydrogenases (*xoxF*) and heterotetrameric methanol dehydrogenase (*mxaFI*) in the *Methylococcales*-affiliated MAGPMV2.70_sub. Only XoxF methanol dehydrogenase was encoded in MAGs affiliated to the classes *Gammaproteobacteria* (PMV6.23), *Alphaproteobacteria* (order *Dongiales*) (PMV8.34_sub) and *Acidobacteriae* (PMV 1.51, PMV 3.11).

#### Methanogenesis

potential was absent in the Movile sediment datasets as suggested by the absence of methyl coenzyme-M reductase (*mcrA*) marker gene across investigated MAGs.

#### CO_2_ fixation

The CO_2_ fixation is presumably critical in a chemoautotrophic ecosystem such as Movile Cave. To verify the autotrophic potential of the sediment communities, the presence of genes for the key enzymes of CO_2_ fixation pathways, namely the Calvin–Benson–Bassham (CBB) cycle (key enzyme: ribulose 1,5-bisphosphate carboxylase/oxygenase (RubisCO), genes *cbbL* and *cbbS*), the reverse tricarboxylic acid (rTCA) cycle (key enzyme: ATP citrate lyase, genes *aclA* and *aclB*) and the reductive acetyl-CoA, or Wood-Ljungdahl (WL) pathway (key enzyme: CO dehydrogenase/acetyl-CoA synthase (CODH/ACS complex), genes *acsA* and *acsB*) was investigated.

No carbon fixation key enzyme was encoded by MAGs from PMV6 and PMV7 datasets. RuBisCO large subunit (*cbbL*) (CBB cycle) was predicted in all other datasets in MAGs affiliated to *Ca*. KSB1 (PMV1.maxbin.001), *Chloroflexota* (PMV2.24), *Micrarchaeota* (PMV4.13) phyla and classes *Alphaproteobacteria (genus Methylocella* (PMV3.41) and order *Dongiales* (PMV8.34_sub)) and *Gammaproteobacteria* (order *Thiohalomonadales*) (PMV4.23). The *Thiohalomonadales* (PMV4.23) and the *Dongiales* (PMV8.34_sub) MAGs encoded both RuBisCO subunits (cbbL/S). Most of the enzymes of the rTCA cycle are shared with the TCA cycle, except for the ATP citrate lyase (ACL) used by autotrophic prokaryotes for conversion of citrate to acetyl-CoA. The rTCA cycle hallmark enzyme subunits were annotated in MAGs affiliated with phyla *Campylobacterota* (PMV4.maxbin.013) and *Nitrospirota* (PMV8.22).

For the anaerobic carbon fixation via WL pathway, the marker genes of CODH/ACS complex, *acsAB*, were found only in PMV4 dataset. The Western branch (carbonyl) was encoded by MAGs affiliated to phyla *Nitrospirota*, (order *Thermodesulfovibrionales*) (PMV 4.maxbin.018) (AcsA (β), AcsB (α), AcsD (δ), AcsC (γ)) and *Ca*. KSB1 (PMV 4.37) (AcsA (β), AcsB (α), AcsC (γ)). The Eastern branch (methyl), for CO_2_ conversion to 5-methyl-tetrahydrofolate, was also encoded in the *Nitrospirota*-affiliated MAG. The *Ca*. KSB1-affiliated MAG encoded for methylenetetrahydrofolate dehydrogenase (*folD*) of WL pathway Eastern branch as well as the acetyl-CoA to acetate enzymes: acetate kinase (*ackA*), phosphate acetyltransferase (*pta*) and a putative phosphotransacetylase. This suggests that the *Ca*. KSB1 MAG could be unable to gain ATP from acetyl-CoA degradation. Enzymes for the Eastern branch (methyl) of WL pathway were encoded in all sediment datasets.

Interestingly, none of the genes of interest for biogeochemical cycling of S and N, CH_4_ oxidation and CO_2_ fixation were annotated in MAGs assigned to *Ca*.Patescibacteria or *Myxococcota* phyla.

All the functional genes implicated in the sulfur and nitrogen cycling, methane oxidation and CO_2_ fixation annotated in MAGs from Movile Cave sediment are listed in Additional file 5: Table S9 to S12 and an overview of the genes, encoding MAGs and pathways are shown in Fig. 4.

### Potential metabolic interactions and dependencies within the communities

To gain a deeper insight into the microbial metabolic webs in Movile Cave sediments, we used the MAGs to construct metagenome-scale metabolic models (metaGEMs) for simulation and prediction of potential metabolic interactions and dependencies within the communities. Noteworthy, no metabolic models could be constructed for three MAGs belonging to *Ca*.Patescibacteria (PMV2.56, 2.61 and 2.72) and thus these were excluded from the analysis. The inability to construct metabolic models for MAGs belonging to *Ca*.Patescibacteria can be a consequence of their reduced genomes size (usually <1 Mb). The rest of the metaGEMs could generate biomass and reproduce growth in the simulation conditions as evidenced by MEMOTE tests results included in the Additional file 6: Table S13. The competition-cooperation potential predicted by SMETANA (Fig. 5A.; Additional file 6: Table S14) was highest in the communities associated with the upper gallery (e.g. PMV7). This was evident especially in the case of the necessary resources overlap (competition). The pattern persisted regardless of metaGEMs reconstruction or communities ‘simulation parameters. When community simulation assumed minimum nutrient availability, as presumed for Movile Cave environment, the community members of the upper gallery (samples PMV6 to 8) exhibited the highest similarities of metabolic requirements (competition) and highest potential for community self-sufficiency (cooperation) (Fig. 5A.b.).

**Figure 5.**
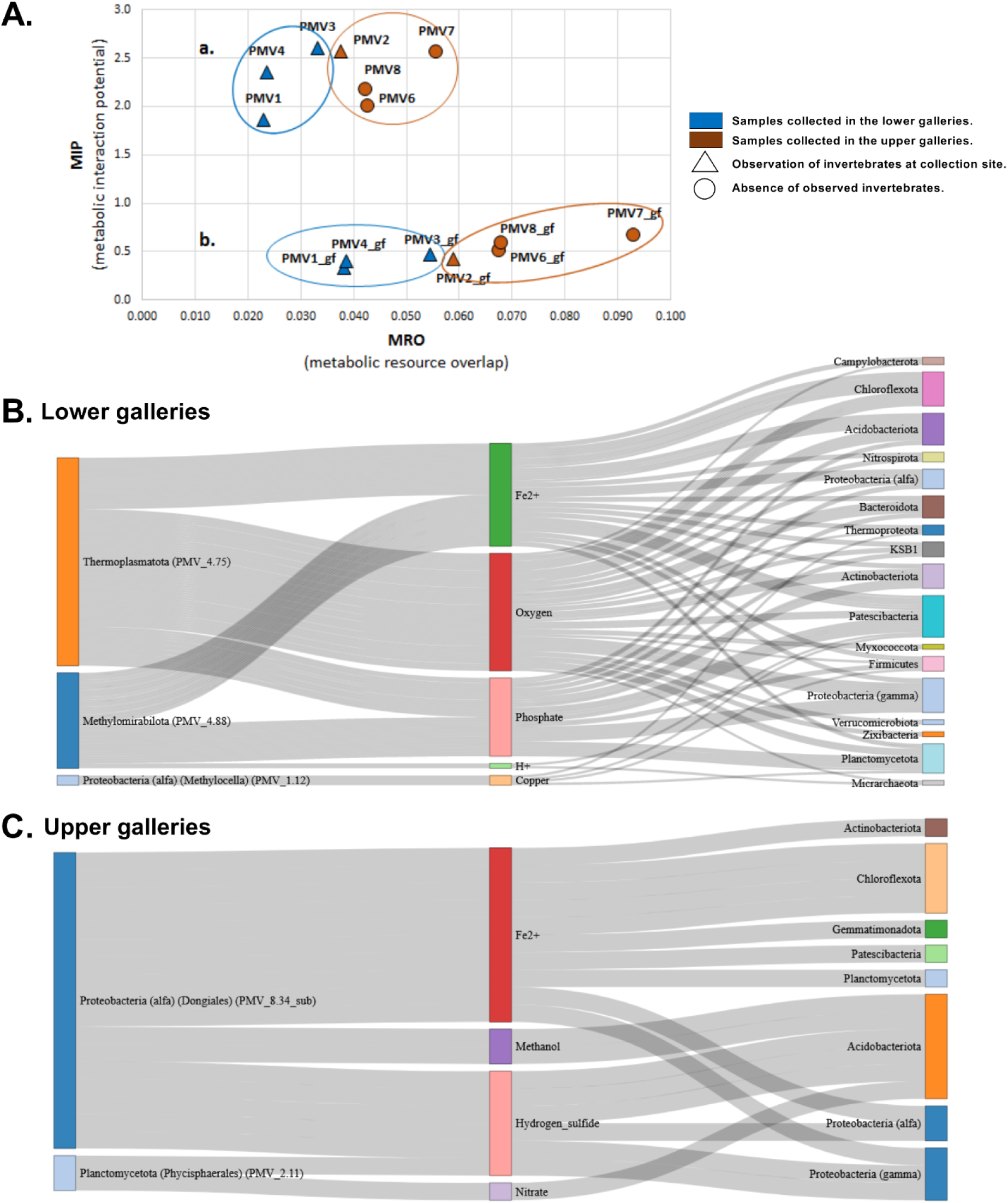
Competition-cooperation landscape of each sample and cross-feeding interactions across wet and dry galleries. **A**. The competition (MRO) and cooperation (MIP) scores (divided by the numbers of MAGs in the community) are shown for different reconstruction and simulation parameters: (**a**.) metaGEMs reconstructed only on genetic evidence and community simulation on complete medium (unconstrained environment); (**b**.) metaGEMs reconstructed by gap-filling for minimal media and community simulation on minimal media (constrained environment). **B**. and **C**. Alluvial diagrams showing compounds exchanged (SMETANA score = 1) in each condition (lower/wet and upper/dry gallery) between the donor (left) and receiver (right) phyla. The colors are used only to distinguish distinct components of the alluvial diagrams.

For an insightful analysis into the microbial metabolic webs in the Movile Cave sediments, we grouped the samples based on distance, namely samples from the lower ‘wet ‘gallery (PMV1, PMV3, PMV4) and samples from the upper dry gallery (PMV2, PMV6, PMV7, PMV8). We examined the cross-feeding interactions and keystone phyla across those two conditions (‘wet ‘and ‘dry’). Our simulation results (SMETANA score = 1) suggested that only ‘currency’ molecules and inorganic ions (O_2_, PO_4_^3-^, H^+^, Fe^2+^, Cu^2+^) were exchanged in the lower gallery. Here, the donor MAGs belonged to archaeal *Ca*.Thermoplasmatota (PMV4.75), bacteria candidate division NC10 (*Methylomirabilota*) (PMV4.88) and the genus *Methylocella* (PMV1.12) (Fig. 5B.). In the upper gallery, besides inorganic ions (Fe^2+^, NO^3-^), hydrogen sulfide (H_2_S) and methanol (CH_4_O) were also exchanged. Here, the essential donors were affiliated to the *Alphaproteobacteria* within order *Dongiales* (PMV8.34_sub) and *Planctomycetota within* order *Phycisphaerales* (PMV2.11) (Fig. 5C.). For both conditions, the taxonomic affiliation of receiver MAGs was diverse, represented by MAGs belonging to 8 and 17 phyla. The MAGs that interact in each simulation condition and their role as donors or recipients of exchanged metabolites are listed in Additional file 6: Table S15 and S16.

Out of the predicted essential compounds (SMETANA score = 1) that are readily exchanged among MAGs, H_2_S, NO^3-^, methanol, and H^+^ exchanges were significantly different (Wilcoxon rank-sum test, BH adjusted p-values < 0.001) across upper and lower galleries (Additional file 6: Table S17).

## Discussions

Our current understanding of microbial life within the Movile Cave ecosystem was limited to the hydrothermal waters, as only microbial mats, water samples and Lake sediments were previously investigated [18, 19]. Our study gives a first insight into the cave’s chemoautotrophic and primary production potential beyond the hydrothermal waters, namely the cave’s sediments.

Based on physicochemical and mineralogical characterization, sampled sediments showed a very different chemical composition, even if located just meters away from one another. PMV6 and PMV8 differed the most from the other samples, followed by PMV3 and PMV4. Mineralogy partially supports such chemical discrepancies, but some of the best represented elements are of different, unknown origin, probably linked to the history of the region where the cave is located and the input of the underground sediments from the Miocene to the more recent climatic events.

The taxonomic novelty of recovered MAGs was high, with over 60% unaffiliated to a genus and 30% classified by RED (Relative Evolutionary Divergence) as new species of established GTDB taxa. The community was diverse with 22 microbial phyla, mainly *Acidobacteriota, Chloroflexota, Proteobacteria, Planctomycetota, Ca*.Patescibacteria, *Thermoproteota, Methylomirabilota*, and *Ca*.Zixibacteria based on the relative abundance of MAGs in samples. In spite of the high novelty at lower taxonomic levels, the overall phyla diversity of sediment MAGs was typical of sulfidic and non-sulfidic cave microbiology [51–53]. Something worth noticing is the lack of *Betaproteobacteria* among recovered MAGs, since it was previously demonstrated for the Movile Cave ecosystem [16, 21].

Functional analyses suggested the presence of chemolithoautotrophic primary producers in the cave sediments from both galleries. As expected, sulfur oxidation, proposed as the ecosystem’s driving force, was mainly present in the lakeside dataset PMV4. The complete oxidation pathways (Sox and Dsr pathways), RuBisCO and partial nitrate/nitrite respiration and denitrification were encoded by the *Thiohalomonadales-affiliated MAG* (class *Gammaproteobacteria*). As *Thiohalomonas* members are obligate chemolithoautotrophic facultative anaerobic sulfur-oxidizing bacteria [54], this MAG might play a key role as primary producer within the ecosystem. It uses reduced sulfur compounds as energy sources, nitrate as an electron acceptor and assimilating CO_2_ via the Calvin-Benson-Bassham cycle. Another primary producer inferred from the Lake edge sample was an *Arcobacteraceae*-related MAG (phylum *Campylobacterota*/*Epsilonproteobacteria)* that encoded the capacity for sulfur oxidation (Sox) potentially coupled with nitrate-reduction (NapB) and CO_2_ fixation via the reverse TCA cycle (*aclAB*). Those are typical metabolic features of *Epsilonproteobacteria* chemolithotrophic primary producers from the deep-sea hydrothermal vent [55]. Similar metabolic traits were found for the recently characterized *Thiovulum* sp. isolated from Movile Cave’s hypoxic Air Bells microbial mat [22]. Moreover, *Arcobacteraceae* family was found dominant in the white filaments from the thermal sulfidic spring of Fetida Cave, Italy [56]. Sulfur oxidation was not however limited to the lakeside sediments. Genes coding for Sox pathway (*soxCD*) were found in PMV2, 6 and 7 datasets of the upper, dry gallery across MAGs affiliated to *Alpha-, Gammaproteobacteria* and *Gemmatimonadota* phylum. The *Gemmatimonadota* role in the sulfur cycle is uncertain, but MAGs with similar attributes were identified in the Siberian soda lakes [6]. In the sediments from around the Lake, sulfur respiration was also evident, which is less critical in self-sustaining ecosystems and a characteristic of heterotrophs. In addition to previous reports [16, 21] describing sulfate reducers in water and microbial mat samples from Movile Cave that were associated to *Deltaproteobacteria*, we assembled a hypothetical thermophilic sulfate reducer MAG affiliated to *Nitrospirota* (order *Thermodesulfovibrionales*) possessing DSR pathway genes (*aprAB*). These results might indicate that sulfate reduction is important in and around the sulfidic water but not away from it.

Starting from the relatively high (0.2–0.3 mM) ammonium concentrations in the cave waters [57] it was formerly implied that nitrogen oxidation might also be a driving force for the ecosystem [21]. A few *Nitrospirota*-affiliated MAGs were identified in the upper and lower galleries, but none of the nitrite or ammonia-oxidation genes were annotated. No nitrifying bacteria were identified based on gene annotation, but an ammonia oxidizing *Nitrososphaera* archaeon encoding for an AmoA-like subunit was assembled from the Lake Room PMV3 sediments. *Nitrososphaera* archaea are facultative chemolithoautotrophs with potential metabolic flexibility [58, 59]. Our metabolic annotation results suggest that the nitrogen cycle throughout the Movile Cave sediments are mostly driven by nitrate/nitrite respiration and denitrification. Those nitrogen reduction processes are linked to taxonomically diverse bacteria and spread among all investigated locations. The dissimilatory and assimilatory nitrate reduction to ammonia was predicted in all datasets and emphasized the ability of the microbial communities to provide bioavailable N for the other trophic links in the Movile Cave ecosystem.

Methanotrophs were also postulated as primary producers in the water and microbial mats [19–21, 24, 60] as 1– 2 % methane concentration was highlighted, especially in the Air-Bells [57]. In this study we were able to assemble methanotrophic MAGs from both lower and upper galleries. These were affiliated to the uncultured UBA1147 genus of *Methylococcales* (in PMV1, PMV2, PMV6) and *Methylocella genus* of *Rhizobiales* (in PMV1, PMV3, PMV6) and encoded for subunits of pMMO monooxygenase. Although no methane-oxidation gene could be annotated, *Methylomirabilota-affiliated* (candidate division NC10) MAGs were found in PMV2 (order *Rokubacteriales*), PMV4 and PMV7 (order *Methylomirabilales*) datasets. Interestingly *Methylomirabilales-affiliated MAG* was one of the most abundant MAGs in PMV7. The presence of *Methylococcales (Methylocccus, Methylomonas, Methylocaldum*) and *Rhizobiales* (*Methylocystis/Methylosinus, Methylocella*) methanotrophs was previously postulated in the microbial matsbased on pMMOA-sequence clones from a CH_4_-enriched culture [19, 20]. The genus *Methylocella* comprises facultative methanotrophs that utilize multicarbon compounds (acetate, pyruvate, succinate, malate, and ethanol) [61]. While the order *Rokubacteriales* might comprise non-methanotrophic bacteria [62], *Methylomirabilales* are known autotrophic (CO_2_ fixing via CBB cycle), denitrifying methanotrophs that can oxidase methane anaerobically. The molecular oxygen needed for methane oxidation is generated by reducing nitrate to dinitrogen gas and O_2_ and bypassing the nitrous oxide formation [63–65]. This is considered the fourth biological pathway known to produce oxygen besides photosynthesis, chlorate respiration, and the detoxification of reactive oxygen species [63].

The key enzymes for CO_2_ fixation were found in 5 out of 7 sediment samples (except PMV6, PMV7), including PMV8 collected farthest from the lake. Noteworthy, in PMV8 MAGs both the CBB and rTCA enzymes were annotated. WL pathway of anaerobic CO_2_ fixation was postulated only in the water edge dataset PMV4. A sulfur-respiring (*aprAB*) MAG affiliated to the order *Thermodesulfovibrionales* (phylum *Nitrospirota*) encoded for the WL pathway key enzymes (*acsAB*). However, the *Thermodesulfovibrionales* order includes members (genus *Thermodesulfovibrio*) that are chemoorganotroph, fermentative and dissimilatory sulfate-reducing bacteria [66]. Therefore, the MAG can use the WL pathway in reverse to break down acetate to CO_2_ and H_2_. Uncultured *Nitrospirota* MAG from PMV8 dataset encoded key genes for rTCA cycle, confirming early evidence revealed by stable-isotope probing (SIP) of water and microbial mat samples that *Nitrospirota* might be able of CO_2_ fixation [21]. CBB cycle key enzymes were detected in most sediment samples, except for PMV6 and PMV7. The archaeal MAG encoding CBB capability found in water edge sample (PMV4) pertains to *Ca*.Thermoplasmatota’s class EX4484-6 that includes MAGs retrieved from marine hydrothermal vent sediments (BioProject PRJNA362212). Thermoplasmata members were also identified in the snottites (thick snot-like biofilm) of Frasassi Cave and gypsum moonmilk of Fetida Cave, both in Italy [51, 67, 68]. The presence of RuBisCO (*cbbL*) and sulfhydrogenase (*hydBG*) subunits in the genome may point towards an autotrophic organism that couples sulfur (S^0^, S_n_) reduction to hydrogen oxidation. Among the bacteria encoding RuBisCO, a MAG affiliated to the order *Dongiales* from the upper gallery PMV8 stands out. GTDB’s *Dongiales* order comprise the members of the formerly known family *Rhodospirillaceae*. The RuBisCO enzyme presence and the lack of genes for the photosynthetic reaction center (*pufLM*) are intriguing for *Rhodospirillaceae*-like bacteria. *Rhodospirillaceae* are known as photoautotrophic, photoheterotrophic and chemoheterotrophic bacteria [69], but not as chemoautotrophs. The *Dongiales*-affiliated MAG may be a chemolithoautotrophic bacteria that fix CO_2_ and uses sulfur compounds as electron sources since the sulfite dehydrogenase (*soeA*) subunit was also annotated in the genome. Another interesting phylum encoding autotrophic features was *Ca*. KSB1 phylum, which currently consists of a single class termed UBA2214. The two *Ca*. KSB1-related MAGs recovered in this study encode oxic- (CBB) and anoxic (WL) CO_2_ fixation potential. Both MAGs also seem to carry nitrate reduction by DNRA and denitrification, which can serve as a nitrogen retainer (DNRA) and a nitrogen remover (denitrification) in the environment.

The metabolic interactions were investigated to extend our understanding of the metabolic potential in sediment-associated microbial communities. Microbial communities in the lower, wet and upper, dry galleries were analyzed using MAG-based metabolic models (metaGEM). Community metabolic modeling approach using metaGEM reconstruction and *in silico* simulation is only recently applied in different fields [70–74]. The community MRO and MIP distribution patterns support the expectation of lower nutrient availability in the dry gallery vs. the wet one. In scant nutrient environments, the microbes compete over the available nutrients (high MRO), and its members might need to have complementary biosynthetic capabilities to decrease their dependency on the scarce external resources (high MIP). In caves, it is known that selfish competition for resources it can be replaced by cooperative and mutualistic associations, such as the ones seen in biofilms [75], maintaining bacterial communities with diverse metabolic pathways, interdependent and cooperative [76]. On the other hand, the MRO/MIP distribution pattern supports the supposition of higher nutrient availability in the areas where invertebrates were present. Those communities (PMV1, PMV2, PMV3, PMV4) had lower MRO, and MIP than the communities where no invertebrates were identified (PMV6, PMV7, PMV8). The simulated cross-feeding interactions highlighted the key donor MAGs for each of the conditions as *Methylomirabilales* and *Methylocella* methanotrophs and *Thermoplasmatota* autotrophic archaeon for the lower, wet gallery; and the autotrophic *Dongiales* and a *Phycisphaerales*-related MAG for the dry gallery. Little could be deduced for the *Phycisphaerales*-affiliated donor MAG. None of the marker genes for sulfur, nitrogen, methane metabolism, or CO_2_ fixation was assembled or annotated. This is not surprising since the phylum *Planctomycetes* has the highest values (35–65 %) of protein sequences with unknown function among bacterial phyla [77]. Moreover, *Phycisphaerales* MAG is the second of its genus (SLJD01). The first was assembled from the surface sediments of a hypersaline soda lake in Siberia (GCA_007135295). As an observation, MAGs belonging to *Ca*.Patescibacteria could not be modeled or were on the receivers’ side of cross-feeding interactions. This is typical for *Ca*. Patescibacteria members with a reduced genomes size (usually < 1.5 Mb) that lack essential biosynthetic capacities, have metabolic dependence and may have a parasitic or symbiotic lifestyle [78–80].

In the absence of light, the chemolithoautotrophic ecosystems are fueled by the oxidation of reduced compounds such as H_2_S, CH_4_, NH_3_, Fe^2+^ and H^+^. The metabolic modeling community simulations pointed to distinct metabolic dependencies of simple compounds in lower and upper galleries. We postulate that the compound accessibility influences the established microbial dependencies in the environment, as H_2_S, NO_3_^-^ (a result of ammonia oxidation), and CH_4_O (a result of methanotrophy) are more available in the lower gallery than in the upper one. Hence the organism’s dependencies on those compounds (i.e., a likelihood of species A growth depending on metabolite X from species B) are established in the upper gallery, where those compounds are lacking in the environment. Similarly, in the case of O_2_, the dependencies appear in the lower gallery where O_2_ concentration is reduced. Ferrous iron (Fe^2+^) dependencies are characteristics for both conditions probably because under oxidizing conditions iron is found mainly in Fe(III) (oxyhydr) oxide minerals (i.e. goethite).

## Conclusions

The present work addressed the diversity, biogeochemical potential and ecological interaction of sediment-associated microbiome located nearby and at distant places from the sulfidic hydrothermal waters feeding the chemoautotrophic-based Movile Cave. Metagenomic-based approaches have indicated that the diversity of the microbiomes detected in Movile Cave sediments spans a wide taxonomic range and is likely to have a high degree of novelty. This study pinpoints that chemolithoautotrophy as an essential metabolic asset in organic carbon-poor Movile Cave environment that is not confined to the high-redox potential hydrothermal water and nearby sediments but to as far as the most distant sample in the dry gallery. This assumption is supported by the recovery of autotrophic MAGs encoding CO_2_ fixation ability via at least three different pathways (CBB, rTCA, WL). Sulfur oxidation was predicted for microbiome detected nearby sulfidic water, whereas ammonia-oxidation might be active in cave sediments in contrast with apparently absent nitrification. Additionally, methanotrophyhas been inferred across all sampled sediments. Therefore, it seems to play a key role in the primary production along the entire length of Movile Cave, not only in the water proximity. Despite simulation simplifications, our modeling approach postulates that nutrient scarcity is the driving force of competition-cooperation patterns across Movile Cave. The metabolic annotation and simulations point towards the autotrophic and methanotrophic MAGs as key donors in the sediments. Cross-feeding interactions can reveal the limiting compounds in the environment and the notable differences between lower and upper galleries.

Our findings point to the potential ecological roles and interactions of the sediment-associated microbiome and add to the previous microbiological investigations focused on the sulfidic waters in the Movile Cave, thus comprehensively expanding our understanding of peculiar chemoautotrophically-based subterranean ecosystems. Nevertheless, prospective direct metabolic quantitative assessments adjoined by multi-omics and isolation and culturing efforts are needed to further unveil the full microbiological picture of this intriguing cave ecosystem.

## Availability of data and materials

Raw metagenomics datasets and metagenome assembled genome derived from this work are publicly available under the NCBI Bioproject PRJNA777757. Metagenomes were deposited to NCBI-SRA with the accession numbers SAMN22875101-SAMN22875107. Metagenome-assembled genomes have been deposited in NCBI BioSample and are available under the accessions SAMN23768506 - SAMN23768611.

## Acknowledgements

We are grateful to Marius Kenesz and Cristian Sitar for providing some of the samples, Mihai Baciu for the access into the cave, Ionuț Cornel Mirea for the cave map editing. We acknowledge Francisco Zorrilla for his assistance in metabolic simulations and Wilcoxon rank-sum test statistical analysis.

## Authors’ contributions

OTM, HLB, IC, GR conceived the study. OTM and LF performed sample collection. OTM, IC, AEL performed metadata analysis. LF performed the mineralogy and AEL performed the geochemistry measurements. IC and DFB performed DNA extraction. GR performed the bioinformatics analysis. IC and GR analyzed the metagenomic data. IC performed the metabolic modeling and community simulation analysis. IC wrote the manuscript and prepared the figures. OTM and HLB provided scientific input to the manuscript. AHV contributed to the final version of the manuscript. All authors read and approved the manuscript.

## Consent for publication

All authors have read and commented on the manuscript and consent to the publication.

## Ethical approval and consent to participate

This article does not contain any studies with human participants or animals performed by any of the authors.

## Supplementary Information

**Additional file 1** (Word Document)**: Figure S1**. The display of sediment samples collected from Movile Cave. Photographic images of the sediments.

**Additional file 2** (Word Document)**: Table S1**. Minimal media composition used in the reconstruction of metaGEMs and community simulations. Minimal compounds (primarily inorganic) that may be in the inorganic environment or can result from microbial metabolic traits (e.g. sulfur and nitrogen oxidation/reduction, CO2 fixation, methanotrophy) previously shown in Movile Cave environment. Compound name and abbreviation are in accordance with the BIGG database (http://bigg.ucsd.edu). **Table S2**. Metagenome scale metabolic models (metaGEMs) reconstruction and community simulations parameters and variants generated and used in this study. **Table S3**. Codes used for CarveMe v. 1.5.1 [41] metabolic model reconstructions (metaGEMs) and SMETANA 1.1.0 [43] community simulations.

**Additional file 3** (.xls)**: Table S4**. The physico-chemical composition of the aqueous fraction of PMV4 sample collected on the water edge in Movile Cave. **Table S5**. Movile Cave sediment samples extended mineralogy.

**Additional file 4** (.xls)**: Table S6**. Main features and statistics of Movile cave sediments-retrieved MAGs. **Table S7**. Taxonomic assignment of MAGs by GTDB-tk v202. **Table S8**. The binning efficiency and the relative abundance of MAGs in each metagenome.

**Additional file 5 (.xls): Tables S9**. The marker genes for sulfur-cycling and the MAGs that encoded them in each sample. **Table S10**. The marker genes for nitrogen-cycle and the MAGs that encoded them in each sample. **Table S11**. The methane oxidation marker genes and the MAGs that encoded them in each sample. **Table S12**. CO_2_ fixation marker genes and the MAGs that encoded them in each sample. Tables S9-12 are the bases for Figure 4 and contain the pathways, KEGG number assignment (KO), gene, sample number and the MAG’s that possess genes from the addressed metabolic pathways.

**Additional file 6 (.xls): Table S13**. MEMOTE [42] test results regarding the potential of gap-fill models (metaGEMs) initialized for the minimal medium to produce biomass (growth). The biomass production potential is indicated by the biomass production values > 0). **Table S14**. The competition (given by MRO score) and cooperation (given by MIP score) potential for the community in each sample, predicted by SMETANA [43]. **Table S15**. The MAGs that interact in SMETANA simulation for the microbial community in the lower galleries. **Table S16**. The MAGs that interact in SMETANA simulation for the microbial community in the upper galleries. **Table S17**. Compounds exchanged, different between upper and lower galleries according to Wilcoxon rank-sum test.

