## Additional file 1. Figure S1 for "Competition-cooperation in the chemoautotrophic ecosystem of Movile Cave – first metagenomic approach on sediments"

| **Supplementary Figure S1** | | |
| --- | --- | --- |
| 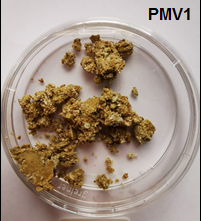 | 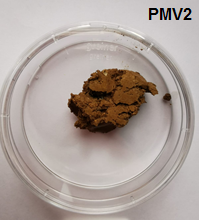 | 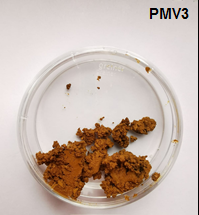 |
| 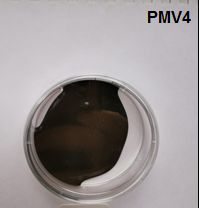 | 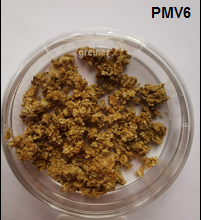 | 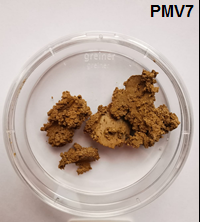 |
| **Supplementary Figure S1.** The display of sediment samples collected from Movile Cave. | | 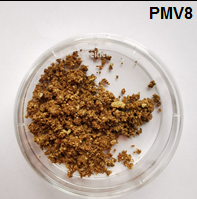 |
