## Additional file 2. Tables S1-S3 for "Competition-cooperation in the chemoautotrophic ecosystem of Movile Cave – first metagenomic approach on sediments"

**Supplemental Tables S1-S3**

**Table S1.** Minimal compounds (primarily inorganic) that may be in the inorganic environment or can result from microbial metabolic traits (e.g. sulfur and nitrogen oxidation/reduction, CO2 fixation, methanotrophy) previously shown in Movile Cave environment. Compound name and abbreviation are in accordance with the BIGG database (<http://bigg.ucsd.edu/>).

| Compound | Abbreviation | Compound | Abbreviation |
| --- | --- | --- | --- |
| O_2_ | o2 | chromite | cro2 |
| H_2_O | h2o | chromate | cro4 |
| Ca^2+^ | ca2 | Pb | pb |
| carbonic acid | h2co3 | Pb_2_ | pb2 |
| bicarbonate | hco3 | uranium (IV) | u4 |
| CO | co | uranium (VI) | u6 |
| CO_2_ | co2 | arsenate | aso4 |
| H^+^ | h | arsenite | aso3 |
| hydroxide ion HO | oh1 | tungstate | tungs |
| Cl^-^ | cl | ammonium | nh4 |
| fluoride | f | nitrogen | n2 |
| Co^2+^ | cobalt2 | ammonia | nh3 |
| Cu^2+^ | cu2 | nitric oxide | no |
| iron | fe | nitrate | no3 |
| Fe^2+^ | fe2 | nitrite | no2 |
| Fe^3+^ | fe3 | nitrous oxide | n2o |
| K^+^ | k | hydroxylamine | ham |
| Mg | mg2 | sulfur | s |
| Mn^2+^ | mn2 | hydrogen sulfide | h2s |
| molybdate | mobd | sulfate | so4 |
| Na^+^ | na1 | sulfite | so3 |
| Ni^2+^ | ni2 | thiosulfate | tsul |
| phosphate | pi | thiosulfate | thios |
| Zn^2+^ | zn2 | trithionate | tton |
| cadmium | cd2 | tetrathionate | tet |
| vanadium (V) | v5 | methan | ch4 |
| vanadium (IV) | v4 | methanol | meoh |
| Hg^2+^ | hg2 | acetate | ac |

**Table S2.** Metagenome scale metabolic models (metaGEMs) reconstruction and community simulations parameters and variants generated and used in this study.

| **PARAMETERS** | | **RESULTS** | |
| --- | --- | --- | --- |
| **Set of reconstructed GEMs**  **(CarveMe)** | **Community simulation (SMETANA)** | **Global interaction**  **within the *sample community***  **(MRO/MIP, competition/cooperation)** | **Detailed cross-feeding**  **dependency *across conditions***  **(upper/lower galleries, Smetana score = 1)** |
| **no_gap_fill**  (only genetic evidence) | **complete media**  (no environmental constraints) | **Yes** | **Yes** |
|  | **minimal media**  (constrained to a minimal environment) | **NO*** | **NO*** |
| **gap_fill**  (to obtain biomass on a  minimal media) | **complete media**  (no environmental constraints) | **Yes** | **Yes** |
|  | **minimal media**  (constrained to a minimal environment) | **Yes** | **Yes** |

*Note*: ***** Models reconstructed only on genetic evidence cannot reproduce growth on minimal media. Results used in this study.

**Table S3.** Codes used for CarveMe v. 1.5.1 (Machado et al., 2018) metabolic model reconstructions (metaGEMs) and SMETANA 1.1.0 (Zelezniak et al., 2015) community simulations.

|  | | **PARAMETERS** | **CODE** |
| --- | --- | --- | --- |
| **GEMs reconstruction (CarveMe)** | | **no_gap_fill** | $ carve MAG.faa --fbc2 -u (grampus/ gramneg/archaea) -o model.xml |
|  |  | **gap_fill** | $ carve MAG.faa --fbc2 -u (grampus/ gramneg/archaea) --gapfill minimal.media --mediadb my.minimal.media .tsv -o model.xml |
| **MEMOTE** | |  | $ carve MAG.faa --fbc2 -u (grampus/ gramneg/archaea) --gapfill minimal.media --init minimal.media --mediadb my.minimal.media .tsv -o model.xml |
| **Community simulations (SMETANA)** | **Global interaction**  **(MRO/MIP)** | **no_gap_fill**  **complete media** | $ smetana *.xml --flavor bigg --molweight --global --debug |
|  |  | **gap_fill**  **minimal media** | $ smetana *.xml --flavor bigg --molweight -m minimal.media --mediadb my.minimal.media.tsv --global --debug |
| **Community simulations (SMETANA)** | **Detailed cross-feeding**  **(Smetana score)** | **gap_fill**  **complete media** | $ smetana *.xml --flavor bigg --molweight --detailed --debug |
|  |  | **gap_fill**  **minimal media** | $ smetana *.xml --flavor bigg --molweight -m minimal.media --mediadb my.minimal.media.tsv --detailed --debug |
| [**https://carveme.readthedocs.io/en/latest/index.html**](https://carveme.readthedocs.io/en/latest/index.html) | | | |
| [**https://smetana.readthedocs.io/en/latest/index.html**](https://smetana.readthedocs.io/en/latest/index.html) | | | |
